# Entrectinib - a SARS-CoV-2 inhibitor in Human Lung Tissue (HLT) cells

**DOI:** 10.1101/2021.09.07.459123

**Authors:** Alejandro Peralta-Garcia, Mariona Torrens-Fontanals, Tomasz Maciej Stepniewski, Judit Grau-Expósito, David Perea, Vikram Ayinampudi, Maria Waldhoer, Mirjam Zimmermann, Maria J. Buzón, Meritxell Genescà, Jana Selent

**Affiliations:** Research Programme on Biomedical Informatics (GRIB), Hospital Del Mar Medical Research Institute (IMIM)—Department of Experimental and Health Sciences, Pompeu Fabra University (UPF), 08003 Barcelona, Spain; InterAx Biotech AG, PARK InnovAARE, 5234 Villigen, Switzerland; Infectious Diseases Department, Vall d’Hebron Institut de Recerca (VHIR), Vall d’Hebron Hospital Universitari, Vall d’Hebron Barcelona Hospital Campus, 08035 Barcelona, Spain

**Keywords:** SARS-CoV-2, COVID-19, specific and non-specific antiviral action, drug repurposing, viral cell entry assays

## Abstract

Since the start of the COVID-19 outbreak, pharmaceutical companies and research groups have focused on the development of vaccines and antiviral drugs against SARS-CoV-2. Here, we apply a drug repurposing strategy to identify potential drug candidates that are able to block the entrance of the virus into human cells. By combining virtual screening with in vitro pseudovirus assays and antiviral assays in Human Lung Tissue (HLT) cells, we identify entrectinib as a promising antiviral drug. We found that part of the antiviral action of entrectinib is mediated by a non-specific mechanism, likely occurring at the viral membrane level. Such a profile could provide entrectinib with protection against the development of drug resistance by emerging SARS-CoV-2 variants.

## 1. Introduction

Severe acute respiratory syndrome coronavirus 2 (SARS-CoV-2) is the pathogen that causes the novel coronavirus disease-2019 (COVID-19). First detected in Wuhan, China, in December 2019, it quickly spread across the country and, by March 11 2020, the World Health Organization (WHO) declared COVID19 a global pandemic, present in almost every country across the globe. Since then, as of August 2021, 213 million people have been infected, counting 4.4 million deaths worldwide [1].

SARS-CoV-2 belongs to the broad family of coronaviruses, a group of RNA viruses that infect mammals and birds. They are enveloped viruses with a positive-sense single-stranded RNA genome and a nucleocapsid of helical symmetry. In humans and birds, they cause respiratory tract infections that can range from mild to lethal. Mild illnesses in humans include some cases of the common cold (which is also caused by other viruses, predominantly rhinoviruses), while more lethal varieties, like SARS-CoV-2, can be fatal.

Major pharmaceutical companies have focused on vaccine development as the primary response to the COVID-19 outbreak. At the same time, several research groups and some pharmaceutical companies are looking for antiviral drugs, targeting viral proteins including the spike protein, the RNA-dependent RNA polymerase (RdRp or nsp12), and the main protease (Mpro or nsp5), to be used as non-immunological interventions. In this context, repurposing of approved drugs represents a promising strategy to find drug candidates at a lower cost and in a shorter time [2]. Currently, several drugs are being used or are under investigation for use against SARS-CoV-2, such as remdesivir (RdRp inhibitor) [3], ritonavir, lopinavir and PF-07321332 (inhibitors of viral proteases) [4–6], or plitidepsin (inhibitors of the replication in the cytoplasm) [7].

However, both vaccines and antivirals targeting virus proteins are at permanent risk of becoming ineffective due to the evolution of viral genomes. Unfortunately, such an evolution is essentially a given. Viruses evolve very quickly, more so when having a high expansion rate such as of the current SARS-CoV-2 pandemic. Many SARS-CoV-2 variants already emerged, as a result of mutation events in the virus genome. Although most mutations are expected to be either deleterious and swiftly purged or relatively neutral, a small proportion can affect functional properties and alter infectivity, disease severity or interactions with host immunity. Indeed, several emerging mutations have been described that show substantially increased infectivity. The WHO has currently declared four variants of concern [8]. The Alpha variant (lineage B.1.1.7) emerged in the United Kingdom in September 2020, with evidence of increased transmissibility and virulence. Notable mutations include N501Y and P681H. An E484K mutation in some lineage B.1.1.7 virions has been noted and is also tracked by various public health agencies. The Beta variant (lineage B.1.351) emerged in South Africa in May 2020, with evidence of increased transmissibility and changes to antigenicity, with some public health officials raising alarms about its impact on the efficacy of some vaccines. Notable mutations include K417N, E484K and N501Y. The Gamma variant (lineage P.1) emerged in Brazil in November 2020, also with evidence of increased transmissibility and virulence, alongside changes to antigenicity. Similar concerns about vaccine efficacy have been raised. Notable mutations also include K417N, E484K and N501Y. Finally, the Delta variant (lineage B.1.617.2) emerged in India in October 2020. There is also evidence of increased transmissibility and changes to antigenicity.

The emergence of SARS-CoV-2 variants with increased transmission rates, severe disease progressions, and/or resistance to vaccinal campaigns and drugs poses a serious threat to global health. In this respect, there is an urgent need to provide effective antiviral drugs with increased resistance to SARS-CoV-2 evolution. In this work, we apply a drug repurposing strategy identifying entrectinib as a potent antiviral drug in vitro which translates into a promising antiviral profile in human lung tissue. Of note, a non-specific antiviral mechanism of action contributes to the overall effect likely occurring at the viral membrane level. Such a profile could provide entrectinib with the ability to be largely resistant to virus evolution and emerging SARS-CoV-2 variants.

## 2. Results

A promising strategy to prevent SARS-CoV-2 infections is to inhibit the initial step of viral entry into the host cell. The SARS-CoV-2 spike protein and in particular the receptor binding domain (RBD) are crucial for viral attachment to the Angiotensin-converting enzyme 2 (ACE2) host cell receptor. In a first step, we explored which binding contacts in the RBD-ACE2 interface are most relevant for viral attachment. This information was exploited in a virtual screen for drug candidates that are able to interfere with the viral attachment interface. Promising candidates were validated in cell-based as well as in human lung tissue assays.

### 2.1 Targeting the RBD-ACE2 interface for a SARS-CoV-2 specific antiviral action

#### 2.1.1 Molecular dynamics simulations of the RBD-ACE2 interface reveal contact hotspots

To identify which residues of the RBD are most relevant for establishing the ACE2 interactions, we explored the RBD-ACE2 interface in long scale molecular dynamics simulations with 10 μs simulation time (Figure 1A and see *Materials and Methods* section). Relevant hotspots were detected by computing the contact frequencies between individual residues. Critical regions with more than 90% contact frequencies were mapped on to the RBD-ACE2 interface and involved the following residues: ARG439, GLN493, GLY496, THR500, ASN501, GLY502 and TYR505 (Figure 1B). They represent valuable information to guide the subsequent virtual screen.

**Figure 1.**
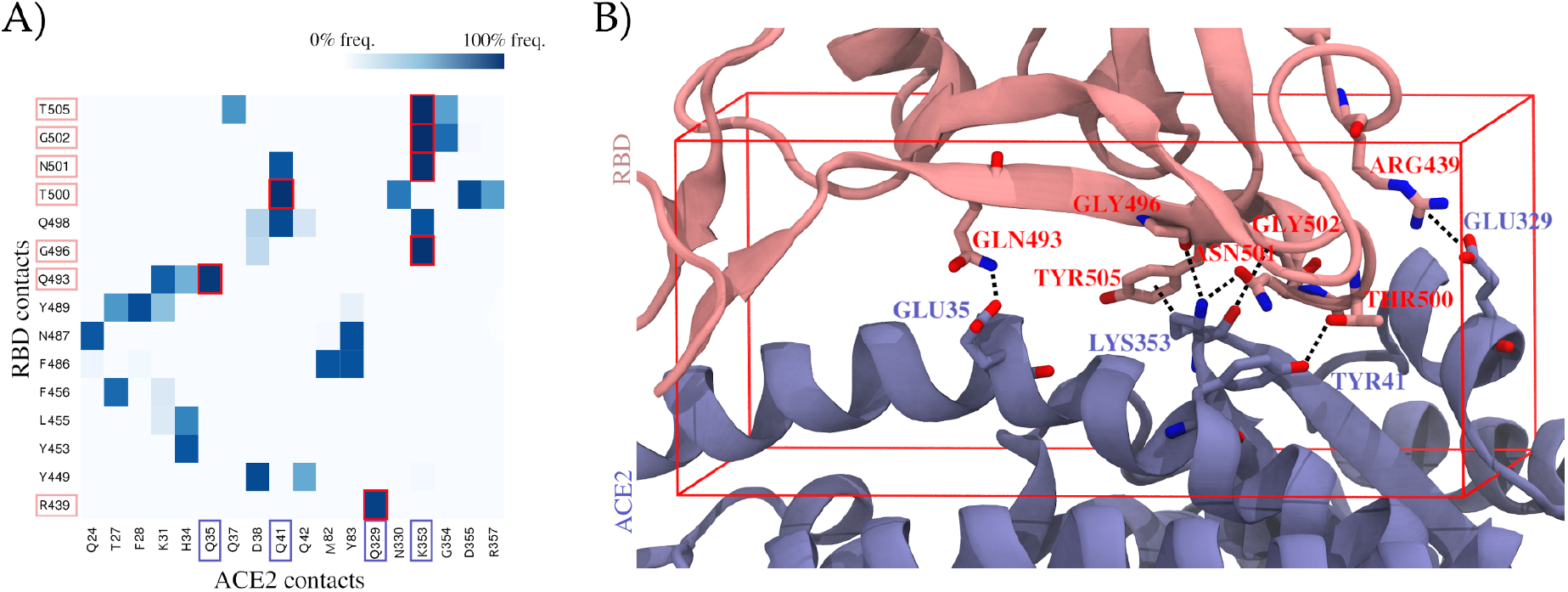
Contribution of individual residues to the stability of the binding complex formed by the spike protein’s receptor binding domain (RBD) and the Angiotensin-converting enzyme 2 (ACE2), computed as contact frequencies. **(A)** Stability of RBD-ACE2 contacts. Heatmap of contact frequencies between RBD and ACE2 residues, where contact frequencies are represented as a color scale from white (0%) to dark blue (100%). In a red square are depicted those contacts which are maintained through > 90% of the simulation. Residues that do not form any interaction with frequency > 50% were filtered out. **(B)** Structural mapping of the most stable contacts (contact frequency > 90%) between RBD and ACE2. The interface region where these contacts are found was used to guide a virtual screening. The docking box applied in virtual screening is indicated in red.

#### 2.1.2 Virtual screening yields two drug candidates with the potential to inhibit SARS-CoV-2 cell entry

For our drug repurposing strategy, we created a curated database of 5,849 compounds including drugs approved by the Food and Drug Administration (FDA) and the European Medicines Agency (EMA), as well as known drug metabolites (see *Materials and Methods* section). The curated database was docked into the RBD-ACE2 interface focusing in particular on regions important for the stability of the complex determined in the previous step (see docking box in Figure 1B and *Materials and Methods* section). We carried out two individual screens targeting the binding interface of the RBD (screen 1) and the ACE2 (screen 2).

Among the top hits of both screenings, we manually selected 5 promising candidates to be tested in a cell-based assay (Table 1). In addition, we included 4 more compounds that had been related to antiviral SARS-CoV-2 activity. (i) Ivermectin, with a proven antiviral effect, was proposed to bind the RBD [9], as well as (ii) argatroban and (iii) otamixaban - two compounds that have been suggested to inhibit viral cell entry in a computational study by interfering with the transmembrane protease TMPRSS2a [10]. Ultimately, we included (iv) apilimod as a positive control - a compound that has been demonstrated to block viral entrance in vitro [2] (Table 1).

**Table 1.**
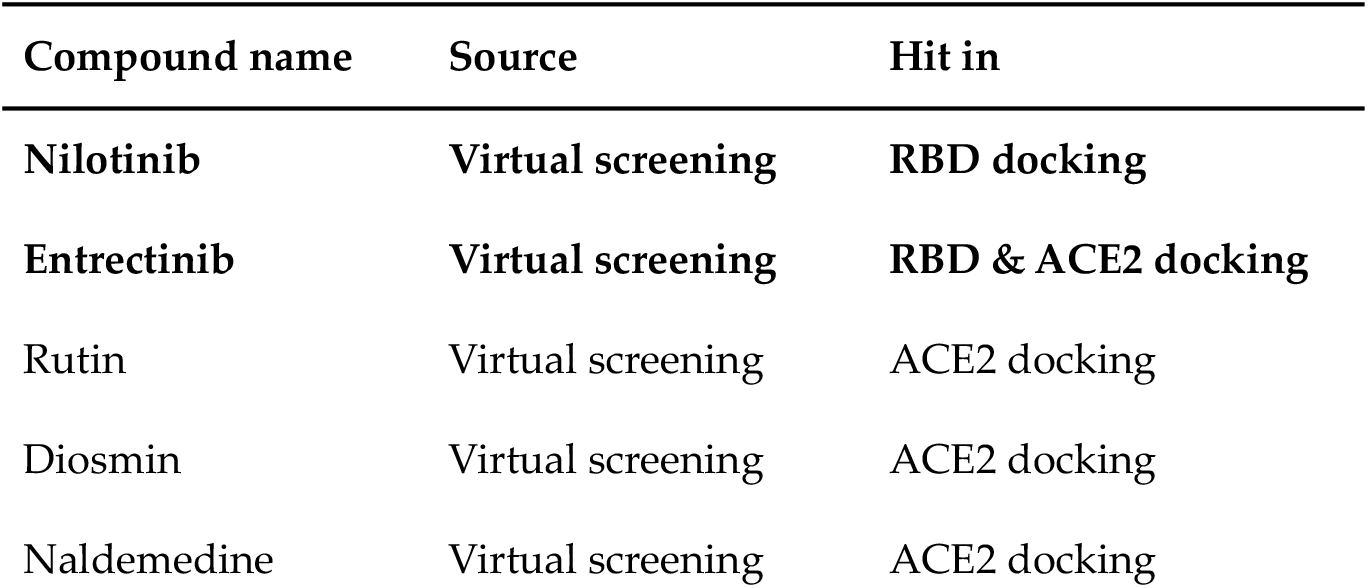
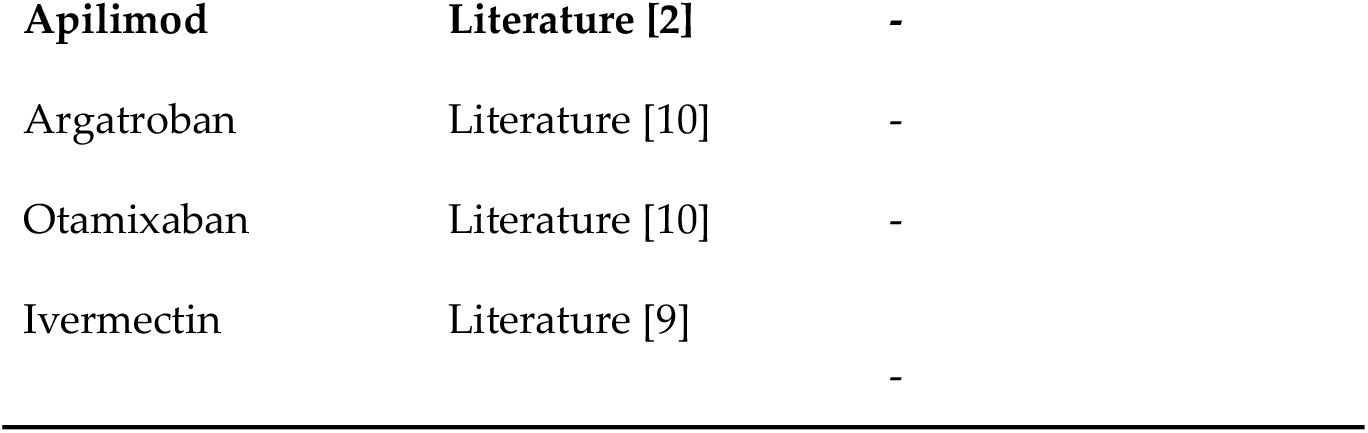
Compounds selected for in vitro inhibition activity assay based on docking or on evidence in literature. Compounds that show an inhibition of virus infection in our in vitro analyses (Figure 2) are highlighted in bold.

**Figure 2.**
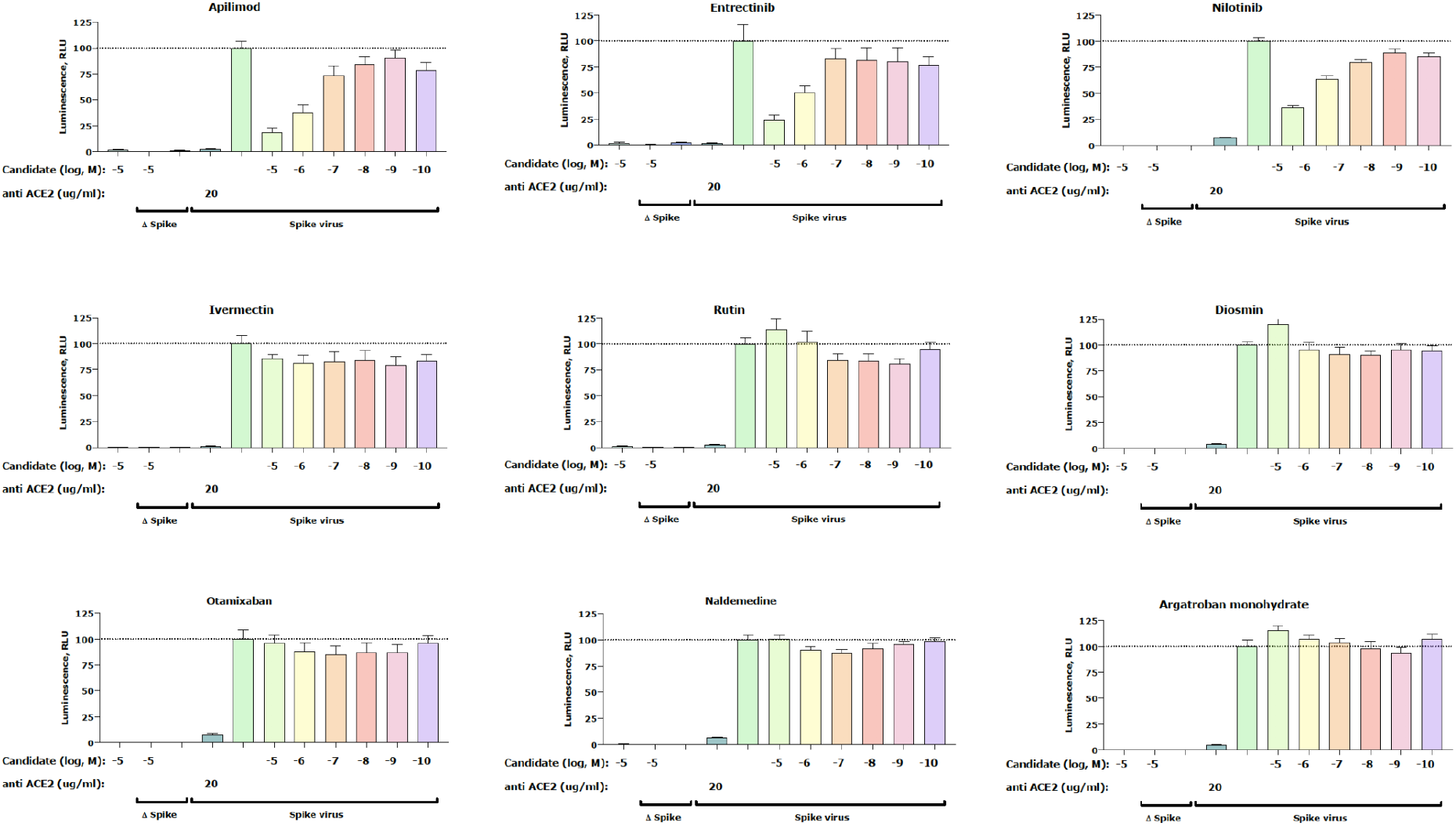
Inhibition of the cell entry of SARS-CoV-2 pseudotyped virus particles. 293T cells transiently expressing human ACE2 and human TMPRSS2a, were pre-treated with the selected compounds at the concentrations indicated for 3 h or with 20 μg / ml of the human anti-ACE2 antibody (positive control), before they were inoculated with SARS-CoV-2-specific spike protein pseudotyped lentivirus particles (spike virus) or particles without a viral envelope (Δ spike). At 48 h postinoculation, pseudotyped virus entry was analyzed by luminescence readout (normalization against untreated spike virus entry). Data represent the mean±SD from three independent experiments carried out in technical triplicates. One biological experiment of otamixaban was excluded due to technical error.

### 2.2 Proof of concept

#### 2.2.1. Pseudovirus assay confirms anti SARS-CoV-2 activity for entrectinib and nilotinib

A total of 9 compounds selected from the virtual screening (5) and literature search (4) (Table 1) were tested in a SARS-CoV-2 pseudovirus assay to measure inhibition of viral cell entry, as described by Walls et al. [11]. For this initial in vitro screen, we tested the inhibitory capacity of selected candidates across 6 different concentrations (Figure 2). Our cell-based assay revealed a dose-dependent inhibitory effect of the positive control apilimod, thus confirming an appropriate experimental setup. In addition to apilimod, we found two more candidates that show a dose-dependent inhibition of virus infection - entrectinib and nilotinib. Interestingly, both of these compounds were predicted to bind with the RBD (Table 1). In contrast, none of the compounds predicted to bind only the ACE2 (rutin, diosmin, naldemedine) showed antiviral activity (Figure 2). Furthermore, no activity was found for argatroban, otamixaban or ivermectin - compounds that had been related to anti SARS-CoV-2 viral activity in the literature [9,10]. Finally, the effect of the isolated compounds on the cell viability was assessed with a standard MTT assay and confirmed that the observed decrease in luciferase signal is not the result of affected cell viability (see *Material and Methods* section).

#### 2.2.2 A non-specific mechanism contributes to the antiviral activity

Compounds that elicit a general and non-specific activity against enveloped viruses (e.g. interference with the viral membrane) can add an advantage against emerging SARS-CoV-2 variants. To interrogate this possibility among the studied compounds, we determined their effect on the cell entry of particles pseudotyped with the membrane protein of vesicular stomatitis virus G (VSV.G) - an enveloped virus lacking the SARS-CoV-2-specific spike protein. Interestingly, we found that apilimod, entrectinib and nilotinib are able to partially inhibit VSV.G infection (Figure 3). This finding suggests that part of the SARS-CoV-2 antiviral action of these compounds does not involve SARS-CoV-2 specific RBD-ACE2 interface and can be attributed to a non-specific antiviral action.

**Figure 3.**
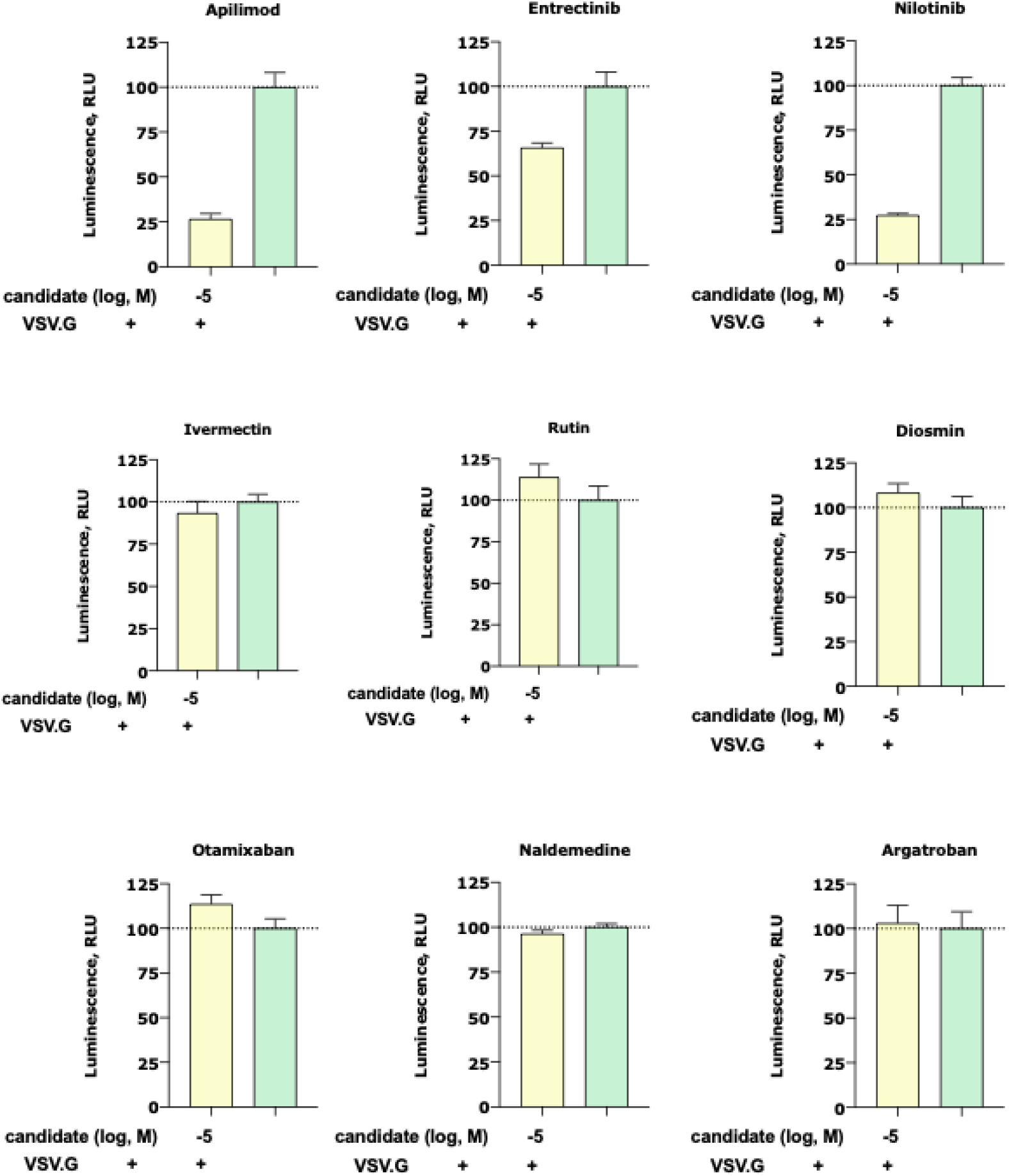
Inhibition of the cell entry of vesicular stomatitis virus G (VSV.G) pseudotyped particles. 293T cells transiently expressing human ACE2 and human TMPRSS2a, were pre-treated with the selected compounds at 10 μM for 3 h before they were inoculated with VSV.G envelope protein pseudotyped lentivirus particles. At 48 h postinoculation, pseudotyped virus entry was analyzed by luminescence readout (normalization against untreated VSV.G virus entry). Data represent the mean±SD from three independent experiments carried out in technical triplicates.

#### 2.2.3. In vitro antiviral activity of entrectinib translates into Human Lung Tissue

To validate if the observed in vitro inhibitory effect of detected candidates (i.e. entrectinib and nilotinib) translates into more native conditions, we carried out an antiviral assay in Human Lung Tissue (HLT) cells [12]. Remarkably, entrectinib exhibited a potent decrease of cellular infection without inducing significant cell death (EC_50_ < 1 μM) (Figure 4). Nilotinib was also found to be active but only at high concentrations (EC_50_ > 15 μM). Interestingly, the positive control apilimod diminished the infection rate to 50% already at very low concentrations, which points to a cytostatic effect of apilimod in this HLT assay. This effect was consistent and confirmed in two repetitive experiments (two different lung donors).

**Figure 4.**
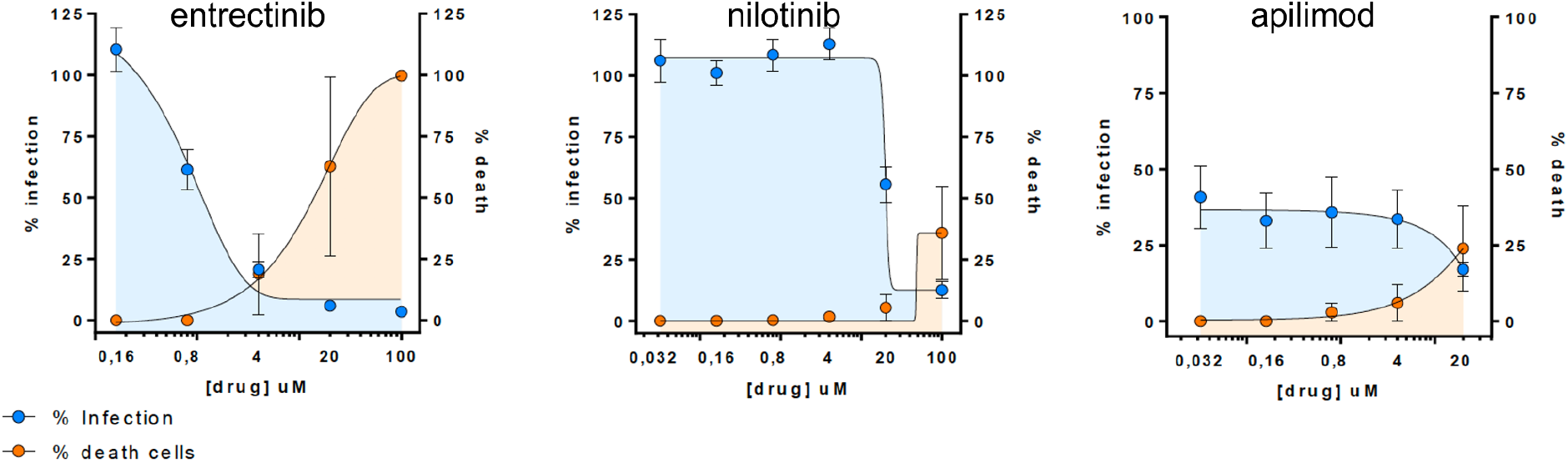
Antiviral assay in Human Lung Tissue (HLT) cells. Percentage of viral entry in HLT cells exposed to VSV*ΔG(Luc)-spike in the presence of the selected compounds. Non-linear fit model with variable response curve from at least two independent experiments in replicates is shown (blue lines). Cytotoxic effect on HTL cells exposed to drug concentrations in the absence of virus is also shown (orange lines).

## 3. Discussion

In this study, we used virtual screening to identify potential drug candidates that are able to block SARS-CoV-2 entrance into human cells by targeting the RBD-ACE2 interface. The most promising candidates were tested for their antiviral activities in a SARS-CoV-2 pseudovirus assay. Thereby, we identified two candidates, entrectinib and nilotinib, that showed similar viral entry blocking effects in vitro compared to the positive control apilimod (Figure 2). Our results for nilotinib and its in vitro antiviral activity against SARS-CoV-2 are in line with a recent study by Cagno et al. [13]. Importantly, the potent antiviral effect of entrectinib observed in our cell-based assay translates into HLT cells at non-cytotoxic concentrations (Figure 4), making entrectinib a promising candidate to combat SARS-CoV-2 infections.

An interesting observation of our study is that entrectinib (an inhibitor of tyrosine receptor kinases A/B/C, ROS1 and anaplastic lymphoma kinase [14]) as well as nilotinib (a Bcr-Abl tyrosine kinase inhibitors [15]) and apilimod (a lipid kinase inhibitor [16]) are able to partially block cell entrance of another enveloped virus that does not belong to the coronavirus family, the VSV.G. This finding indicates that part of the antiviral activity is mediated by an effect that is not specific to SARS-CoV-2. It is tempting to speculate that the mechanism that underlies this non-specific inhibition of viral cell entry occurs at the level of (i) virus-cell fusion or (ii) viral inactivation prior to virus-cell fusion. In fact, tyrosine kinase inhibitors similar to nilotinib have been reported to inhibit virus-cell fusion for SARS-CoV and MERS-CoV in vitro [17,18]. The proposed mechanism involves blocking the Abelson (Abl) kinases Abl1 and Abl2, which are likely involved in coronavirus infection. However, this proposed mechanism would only justify the effect of nilotinib, as entrectinib and apilimod act on different types of kinases. On the other hand, viral inactivation before virus-cell fusion has been proposed for some broad spectrum antiviral agents such as LJ-001 [19]. The antiviral mechanism is based on its ability to intercalate into the viral membrane, which disturbs the virus function. Such mechanism is highly non-specific as the viral envelope (typically derived from the host cell) is largely conserved between enveloped viruses. It is worth noting that such a non-specific mechanism could be beneficial when fighting emerging SARS-CoV-2 variants evading antiviral treatments, such as those based on specific molecular recognition patterns (e.g. vaccines targeting the spike RBD, SARS-CoV-2 specific protease inhibitors).

In conclusion, our study reports a promising antiviral activity of entrectinib against SARS-CoV-2 in human lung tissue, not previously reported. Entrectinib is an FDA-approved drug for the treatment of solid tumors with NTRK fusion proteins and for ROS1-positive non-small cell lung cancers. We find that part of its antiviral action is mediated by a non-specific effect, which likely provides entrectinib with a broad spectrum profile against enveloped viruses. Such a non-specific component can be highly beneficial and protect against the development of drug resistance as a consequence of virus evolution. Future studies have to address the mechanism of action and to confirm these promising data in clinical studies.

## 4. Materials and Methods

### 4.1 RBD-ACE2 interface examination

To find relevant regions that mediate contacts in the interface between the RBD and the ACE2, a 10 μs simulation of the RBD-ACE2 complex (PDB ID 6VW1) from D.E. Shaw laboratory was analysed (Simulation ID: DESRES-ANTON-10875755) [20]. For that, we used the GetContacts 2.0 software [21] to obtain the frequency of total contacts that each interface residue makes during the simulation. Residues with more than 90% contact frequency were considered to be contact hotspots, of relevance for virtual screening.

### 4.2 Database creation

A database of every FDA- and EMA-approved drug was created for the subsequent analyses. For this purpose, DrugBank [22] and PubChem [23] databases were used, merging both of them in a single database in SDF format. Redundancies in both databases were avoided by identifying common IDs. Intermediate drug metabolites were also included from the ZINC15 HMDB Drug Metabolites database [24]. The resulting database included a total of 5,849 compounds.

Next, we removed compounds that could not be properly docked by the molecular docking software (AutoDock Vina 1.1.2 [25,26]), i.e. > 32 rotatable bonds as recommended by the software documentation. Then, different conformations for each compound were found using LigPrep software [27]. Finally, every conformation was converted to PDBQT format using OpenBabel 2.4.0 software [28] (Figure 5).

**Figure 5.**
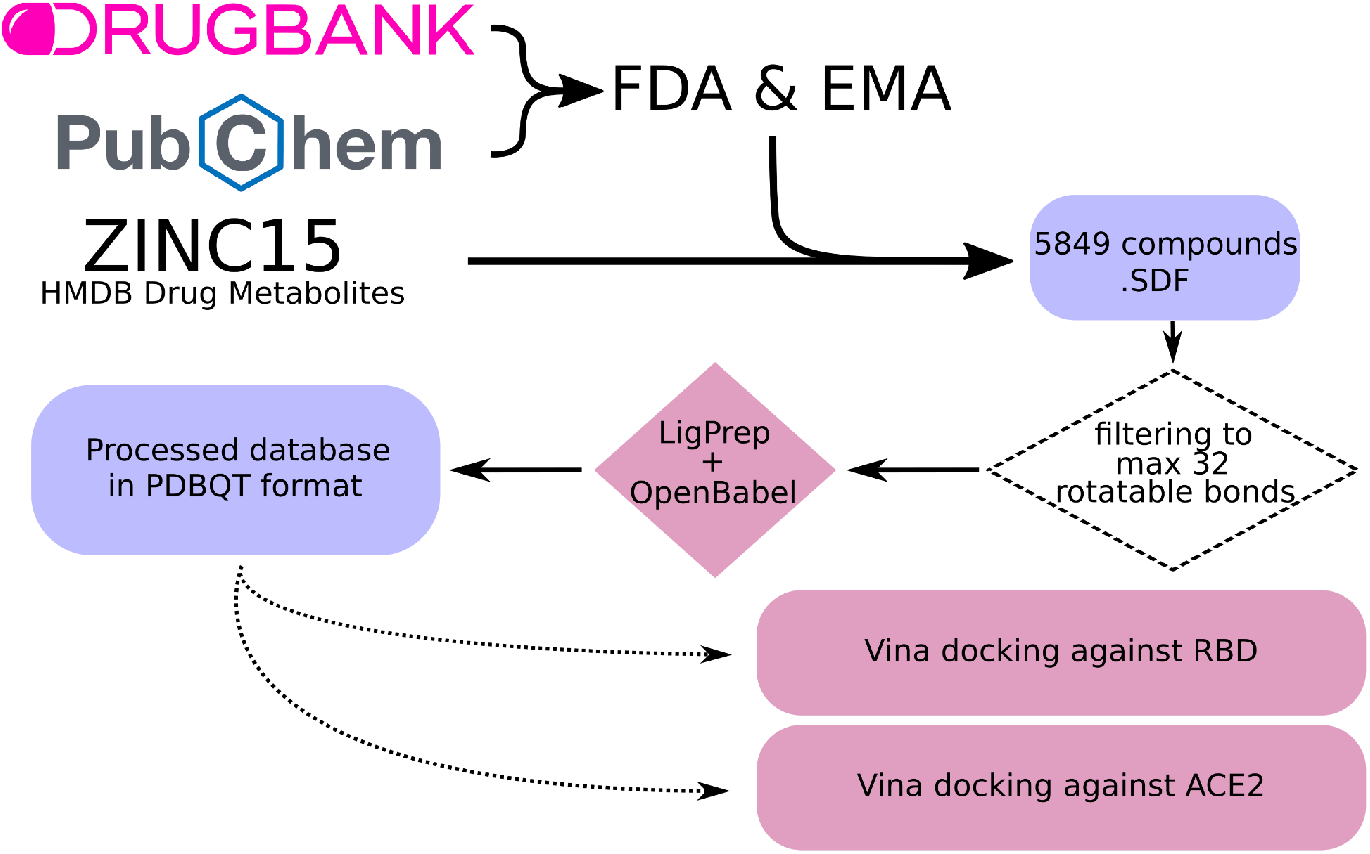
Workflow for the creation of a curated database including drugs approved by the Food and Drug Administration (FDA) and the European Medicines Agency (EMA), as well as known drug metabolites. First, compounds were filtered to discard those with more than 32 rotatable bonds, which are not suitable for docking. Next, different conformations were created using LigPrep [27]. Last, conformations were converted into PDBQT format using OpenBabel [28]. The obtained compounds were used for docking against the RBD and ACE2.

### 4.3 Molecular Docking

For the molecular docking, AutoDock Vina 1.1.2 software [25,26] was used. First, RBD and ACE2 proteins were prepared and converted into PDBQT format using AutoDock Tools 1.5.6 [29]. Both proteins were obtained, in an unbound state, from a simulation from D.E. Shaw laboratory (Simulation ID: DESRES-ANTON-10895671 [20], based on PDB ID 6VW1, replicate 000151, first frame). Then, the docking was performed against the RBD and ACE2, using the previously created database (see *4.2 Database creation* section). The docking region was set to cover the interface contact hotspots detected from the molecular dynamics simulation (see *4.1 RBD-ACE2 interface examination* section), with its dimensions being 18Å × 30Å × 40Å (see docking box in Figure 1B). This process was made using an exhaustiveness parameter of 10. Top hits were analyzed using VMD 1.9.3 software [30].

### 4.4. Plasmids & Cell lines

The Lenti X 293T cell line (632180) was purchased from Takara Bio. Third generation lentivirus packaging plasmids - pLenti CMV Puro Luc (17477), pMDLg.pRRE (12251), pRSV.REV (12253), pMD2.G (12259) were purchased from Addgene. Human ACE2 plasmid was ordered from Genescript. Human TMPRS2 (pUNO1-hTMPRSS2a) was ordered from InvivoGen. Full length SARS-CoV-2 Spike plasmid (VG40589-UT) was purchased from Sino Biological. The Spike ORF was subcloned into pcDNA3.1(+) vector via Gibson Assembly (NEB). Subsequently, a truncated version without the C terminal 21 amino acids, the Spike_CTR plasmid, was generated via Q5 Site Directed Mutagenesis (NEB).

### 4.5. Small molecules

Small molecules were purchased from MedChemExpress at 10mM in DMSO.

### 4.6. Pseudovirus production

On Day 1, Lenti X 293T cells were seeded at a density of 60,000 cells/cm^2^ per T175 flask in 34ml DMEM supplemented with 10% FBS and 1mM Sodium Pyruvate. On Day 2, plasmid co-transfections were performed with 1mg/ml PEI MAX (24765, Polysciences) with DNA:PEI at a ratio of 1:3 as follows. Spike: 20μg pLenti Luc, 20μg pMDLg.pRRE, 9.5μg pRSV.REV & 10.5μg Spike_CTR at a molar ratio of 1:1:1:0.5 respectively. VSV.G: 20μg pLenti Luc, 20μg pMDLg.pRRE, 9.5μg pRSV.REV & 6.5μg pMD2.G at a molar ratio of 1:1:1:0.5 respectively. Δ Spike: 20μg pLenti Luc, 20μg pMDLg.pRRE, 9.5μg pRSV.REV at a molar ratio of 1:1:1 respectively. Plasmids and PEI were prepared separately in 1ml OptiMEM, then combined & incubated at room temperature for 15 min. 2ml DNA/PEI complex was added to the media and mixed gently. On Day 5, media was collected and sterile filtered into 50ml tube via 0.2μm syringe filter. 5ml of 3M NaCl and 10ml of 50% PEG 8000 (MD2-250-13, Molecular Dimensions) was added to the virus supernatant for a final 0.3M NaCl and 10% PEG respectively. Virus was mixed gently by inverting the tube a few times and incubated at 4C for 24 hours. On Day 6, tubes were centrifuged at 4C for 45min at 1500g and the resulting pellet was resuspended in 3.5ml of DMEM + 10% FBS + 1mM Sodium Pyruvate. Aliquots of 1.8ml virus were stored at −80°C. All virus and assay work was performed in a BSL2 facility. Firefly Luciferase, encoded by pLenti Luc was used as the assay reporter. VSV.G, encoded by pMD2.G was used as positive control. Δ Spike was used as negative control.

### 4.7. Pseudovirus assay

On Day 1, 293T cells were seeded at 30,000 per well in white solid bottom 96 well plates that were either Poly L Lysine or Collagen coated in completed media (DMEM + 10% FBS + 1mM Sodium Pyruvate) at 37°C and 5% CO_2_. On Day 2, cells were transfected with 150ng of hACE2 + 15ng hTMPRSS2a per well with Lipofectamine 2000 in OptiMEM according to manufacturer instructions. 6-8 hrs post transfection, the transfection media was replaced with 100μl DMEM + 10% FBS. On Day 3, cells were incubated with 20ul of 6x concentrations in HBSS of either the small molecules/peptides or an anti ACE2 antibody (AG-20A-0037PF-C500, Adipogen) for three hours. Subsequently, frozen virus aliquots were thawed and 100ul of virus solution was added to the respective wells. On Day 5, the assay plates were equilibrated to room temperature, the cells were washed once with 200μl sterile PBS and incubated with 100μl of substrate (50μl PBS + 50μl ONE-Glo EX substrate, E8130-Promega) in the dark for 10min. The luciferase signal was measured on a PHERAstar plate reader (BMG Labtech).

Cell viability was assessed using the CellTiter 96 Aqueous One Solution (G3582, Promega) according to the manufacturer’s instructions. In brief, cells were incubated in 20ul of the CellTiter solution and incubated for 4 hours before measuring absorbance at 490nm on a Flex Station plate reader (Molecular Devices).

All data were analyzed using the GraphPad Prism software 8.

### 4.8 Cell viability assay

Cell viability was assessed as follows. On Day 1, 293T cells were seeded at 30,000 per well in black, clear bottom 96 well plates that are either Poly L Lysine or Collagen coated in cell culture medium (DMEM + 10% FBS + 1mM Sodium Pyruvate). The plates were transferred to an incubator at 37°C and 5% CO_2_ for 18 – 22hrs. On Day 2, 150ng hACE2 + 15ng hTMPRSS2a were transfected per well with Lipofectamine 2000 according to manufacturer’s instructions. 6-8 hrs post transfection, the media was replaced in all wells with 100μl complete media and the plates were returned to the incubator. On Day 3, 20μl of 6x stock of small molecules was dispensed to the respective wells to achieve 1x final dilution. On Day 5, 20μl of CellTiter 96 Aqueous One Solution (G3582, Promega) was dispensed per well. The plates were returned to the incubator for 4hrs before plates were sealed and absorbance at 490nm was measured on Flex Station plate reader (Molecular Devices). Data was analysed on GraphPad Prism software.

### 4.9 Antiviral assays in Human Lung Tissue (HLT) cellsAntiviral assays in HLT cells

The antiviral assays were performed as recently described [12]. Briefly, non-neoplastic areas of lung tissues were obtained from patients undergoing thoracic surgical resection at the Thoracic Surgery Service of the Vall d’Hebron University Hospital. Study protocol was approved by the Ethical Committee (Institutional Review Board number PR(AG)212/2020). Tissue was enzymatically digested with 5 mg/ml collagenase IV (Gibco) and 100 μg/ml of DNase I (Roche) for 30 min at 37°C and 400 rpm and mechanically digested with a pestle. The resulting cellular suspension was washed twice with PBS and resuspended in fresh medium (RPMI 1640 supplemented with 5% FBS, 100 U/ml penicillin, and 100 μg/ml streptomycin) and DNase I to dissolve cell aggregates. Cell number and viability were assessed with LUNA™ Automated Cell Counter (Logos Biosystems). Triplicates of five-fold serial dilutions of the antiviral compounds were tested in HLT cells using 2-3 different donors. Drug dilutions were prepared in R10 in a 96-well plates. HLT cells were added at a density of 300,000 cells/well and incubated with the compound for at least 1h before infection. Then, multiplicity of infection (MOI) 0.1 of VSV*ΔG(Luc)-S virus, generated as previously described with the mutation D614G and a deletion in the last 19 amino acids in the spike (plasmid kindly provided by Dr. Javier García-Pérez, Instituto de Salut Carlos III, Spain) were added to the plates and spinoculated at 1,200g and 37°C for 2h. Cells were then cultured overnight at 37°C in a 5% CO_2_ incubator. Subsequently, cells were incubated with Britelite plus reagent (Britelite plus kit; PerkinElmer) and transferred to an opaque black plate. Luminescence was immediately recorded by a luminescence plate reader (LUMIstar Omega). To evaluate cytotoxicity, we used the CellTiter-Glo® Luminescent kit (Promega), following the manufacturer’s instructions. Data was normalized to the mock-infected control, after which EC_50_ and CC_50_ values were calculated using Graph-Pad Prism 7.

## Author Contributions

Conceptualization. J.S., M.W., M.Z, M.J.B and M.G.; supervision, J.S.; formal analysis, A.P.G, M.T.F., T.M.S., J.G.E., D.P., V.A., M.W., M.Z., M.J.B and, M.G.; writing – original draft, J.S., M.T.F., A.P.G. and T.M.S.; writing – review & editing, J.S., M.T.F., T.M.S., A.P.G, J.G.E., D.P., V.A., M.W., M.Z., M.J.B and, M.G.

## Institutional Review Board Statement

The study was conducted according to the guidelines of the Declaration of Helsinki, and approved by the Ethics Committee of the Vall d’Hebron University Hospital (Institutional Review Board number PR(AG)212/2020).

## Acknowledgements

We wish to thank Alessandro Potenza (InterAx Biotech AG) for support in setting up and performing the pseudovirus assay.

